# Ovarian cancer cells exhibit diverse migration strategies on stiff collagenous substrata

**DOI:** 10.1101/2024.07.08.602474

**Authors:** Madhumitha S, Ramray Bhat

## Abstract

In homoeostasis, the shape of epithelial cells and their sessility, i.e. lack of movement are intricately linked together. Alterations in this relationship as a result of malignant transformation and variations, thereof by the mechanical microenvironments that cancer cells encounter as they migrate are as yet ill-understood. Here we explore the interdependency of such traits in two morphologically distinct but invasive ovarian cancer cell lines (OVCAR-3 and SK-OV-3) under mechanically variant contexts. To do so, we came up with a minimal metric toolkit consisting of velocity, local and global turning angles, persistence of migration, major axis dynamics, morphomigrational angle, and elongation dynamics, and rigorously measured their variation using a Shannon entropic distribution to map heterogeneity in behavior between, and across, trajectories of migration. Two stiffness conditions on polymerized Collagen I with Young’s moduli of 0.5 (soft) and 20 (stiff) kPa were chosen. Both the epithelioid OVCAR-3 and mesenchymal SK-OV-3 cells on soft substrata migrated slowly and in an undirected manner. On stiff substrata, SK-OV-3 showed a persistent directed motion with higher velocity. Surprisingly, OVCAR-3 cells on higher stiffness moved at a velocity higher than SK-OV-3 cells and showed a distinct angular motion. The polarity of SK-OV-3 cells on stiff substrata was well-correlated with their movement, whereas for OVCAR-3, we observed an unusual ‘slip’ behavior, where the axes of cell shape and movement were poorly correlated. An examination of their deformability showed that OVCAR-3 and SKOV-3 on softer substrata were relatively rigid but showed greater shape variation (especially OVCAR-3) on stiffer substrata. Therefore, a rigorous quantification using the above metric toolkit reveals how an interplay between intrinsic deformability and the mechanical microenvironment on pathotypic matrix substrata allow distinct migratory dynamics of epithelioid and mesenchymal ovarian cancer cells.

## INTRODUCTION

Cell migration is an essential phenomenon in organogenesis, homeostasis, wound healing, and immunosurveillance (1). It involves a continuous coordination between physicochemical cues external to the cell and its internal signaling cascades through active cytoskeletal rearrangement (2,3). Abnormal migratory behaviors due to aberrant signaling or changes in the external microenvironment, in turn, have been shown to be strongly associated with histopathological states such as cancer, which is typified by ectopic localization of cells due to their invasion outside native niches: a process referred to as metastasis (4,5). Such changes in the microenvironment have also been seen to have a broad effect on cellular morphology. A well-known correlation between morphology and migration is seen when polygonal epithelial cells transit to a dysmorphic ‘mesenchymal’ state with a concomitant increase in motility (epithelial-to-mesenchymal transition, EMT) (6). Recent studies also show that cancer cells adopt an amoeboid transition state by reducing cell-cell and cell-ECM adhesion and further increase their rate of migration. This constant switch between epithelial to mesenchymal and amoeboid has been postulated to play a major role in cancer cell survival and metastasis (7,8). Current studies have highlighted the importance of measuring cell morphological features as they help incidate the physiological status of a cell and distinguish cancer cells from their untransformed counterparts (9,10). The most common mathematical metrics used currently for quantifying cellular shape include aspect ratio, eccentricity, shape index, and circularity, while for cell migration, it includes accumulated distance, average velocity, turning angles, and mean square displacement (11,12). Advancement in computational approaches has resulted in several machine learning and neural network-based approaches for cellular segmentation and classification models for distinguishing and characterizing various morphology and migratory modes (13–15). However, the complexity of such models draws researchers back to using static mathematical parameters to quantify morphology and migration. As studies look into the characteristics of morphology and migration of cells as separate entities, there is a need to rigorously investigate how correlated these processes are under different microenvironmental contexts (14,16) (see also a recent study by Kolodziej and coworkers which provides fresh insights on the interdependency between both metrics using numerical parameters that incorporate cell geometry, cell orientation, and their corresponding migratory behavior (17)).

The need for decoding cellular morpho-migratory dynamics is particularly relevant to invasive cancer cells, which exhibit distinct morphological features and migratory characteristics. This study focuses specifically on epithelial ovarian cancer, which ranks as the seventh most common cancer in terms of incidence in women as well as the eighth most common cause of death from cancer in women (18). The current study uses one cell line each representing high-grade serous ovarian cancer and non-serous cancer. Although having distinct histopathological features, both cells are known to metastasize in animal models (19).

The ability of cancer cells to metastasize rapidly comes from their ability to change their surrounding microenvironment as well as to adapt to such altering niches (20). Biophysical cues from the extracellular matrix (ECM) microenvironment include stiffness, porosity and topography of the underlying substrate (21,22). Ovarian cancer progression involves cells navigating transcoelomic tissues such as lymph nodes, peritoneal linings, and fibrosed areas that show a range of stiffness from 0.5 to 25 kPa. Hence, in the current study, we investigate the role of substrate stiffness using collagen-coated polyacrylamide hydrogels mimicking normal (soft – 0.5 kPa) and tumor (stiff – 20 kPa) tissue on the spatiotemporal cellular plasticity in ovarian cancer cells. While the motility of transformed mesenchymal cells and that of untransformed epithelial sheets have received extensive attention, behaviors of epithelioid cancer cells within, or on the surface of, ECM are as yet ill-understood.

In this study, we develop a toolkit to rigorously analyze parameters relating to migration and morphology of cancer cells. We then utilize the toolkit to study the role of biophysical cues on ovarian cancer cells’ phenotype over a short but finely sampled spatiotemporal scale. Our study reveals surprising effects of the mechanical microenvironment on the locomotive plasticity of ovarian cancer cells.

## MATERIALS AND METHODS

### 1. Cell culture and time-lapse set up

The current study used two different ovarian cancer cell types – OVCAR-3 and SK-OV-3 (ATCC). The OVCAR3 cells were cultured in RPMI media supplemented with 20% FBS (Gibco), while SK-OV-3 cells (WT and GFP-labelled) were cultured in McCoy’s media supplemented with 10% FBS. Approximately 12,000 cells were seeded per well in an 8-well chamber slide coated with the required substrate. Cells were then kept in a cell incubator with 5% CO_2_ 37°C and humidification for overnight adhesion. The main objective was to obtain sparsely distributed cells to collect single-cell data. Time-lapse experiments were performed on the Inverted Epifluorescence Microscope (IX83 Olympus) at 20X objective for 3 hours. Since, cellular protrusion dynamics are in the range of 120-180 s, snapshots are taken at a time interval of 2 minutes (23,24).

### 2. Polyacrylamide (PA) Gel Preparation

PA gels of required stiffness was prepared in reference to the relative concentrations stated in Engler et. al (25)

#### (a) Activation of glass slide

Activation of glass slides (8-well chamber slides) was performed using 10% (3-Aminopropyl) trioethoxysilane with an incubation time of 20 minutes. Excess silane was removed by washing twice with autoclaved filtered water. Fixation was then done using 0.5% glutaraldehyde for 45 minutes. Excess glutaraldehyde was removed by washing thrice with autoclaved filtered water.

#### (b) Preparation of sandwich coverslip

The sandwich coverslip was coated with Rain-X to provide a hydrophobic coating. After 10 minutes, the excess coating was removed by rinsing twice with autoclaved filtered water.

#### (c) Preparation of hydrogel(s)

A solution of acrylamide and bis-acrylamide of the required concentration for specific stiffness was prepared in autoclaved filtered water. For 0.5 kPa, 0.75 ml of 40% acrylamide and 0.3 ml of 2% bis-acrylamide were mixed in 8.95 ml of autoclaved filtered water. For 20kPa, 2ml of 40% acrylamide and 1.32 ml of 2% bis-acrylamide were mixed in 6.68 ml of autoclaved filtered water. 60 – 80μl of polymerizing gel solution is usually added per well. Therefore, to the required volume of acrylamide and bis-acrylamide solution, 1/100^th^ of the total volume of 10% ammonium persulfate (APS) and 1/1000^th^ of the total volume of tetramethylethylenediamine (TEMED) were added to produce polymerizing gel. This polymerizing gel was immediately added to the activated glass slide. The hydrophobic sandwich coverslip was placed immediately on top of the solution to uniformly spread the gel in the glass slide. The coverslip is usually removed within 5 minutes, as the gel usually forms by that time.

#### (d) ECM coating the gel surface

The gel is initially coated with sulfo-SANPAH, a heterobifunctional protein cross-linker for ECM coating. Since sulfo-SANPAH is a photoreactive cross-linker, the coated gel is exposed to UV for 25 mins. The gel is then washed with phosphate buffer solution (PBS) thrice to remove excess sulfo-SANPAH. To the activated gels, 100μg/ml of collagen was added. An initial collagen (Gibco™) concentration of 1mg/ml was prepared. Initial pre-cooling of all the gel components (type I collagen, 10X DMEM, 2N NaOH, PBS) and an eppendorf tube was done. To prepare a volume of 100 μl of 1mg/ml collagen, 33.3 μl collagen of stock concentration 3 mg/ml was neutralized with 0.67 μl of 2 N NaOH. The solutions were thoroughly mixed until a pink color was observed. To this, 10 μl of 10X DMEM and 56 μl of PBS were added. This was further diluted to 100μg/ml by the addition of PBS. The solutions were thoroughly mixed, and a volume of 100μl was added to each well within an 8-well chamber. The 8 well-chambers are kept in the incubator at 37°C for 45 – 60 minutes for fast polymerization and then transferred into the refrigerator at 4°C overnight to enhance the duration for attachment of collagen protein to the gel surface. The well-chambers are again placed in the incubator 1 hour prior to seeding cells for the experiment.

### 3. Image Processing and Analysis

#### a. Segmentation, binarization, and analysis

The image processing was done in an open-source package, namely, Fiji (26). The raw time-lapse image sequence was converted to .tif format and cropped to obtain fields with a higher number of single cells. Manual thresholding was performed to segment fluorescently labelled cells (GFP-labelled), and get binarized images. To the obtained processed file, a fitted ellipse was overlayed to get the required parameters using an ImageJ Macro code. Cell measurements such as centroid, major axis length, minor axis length, major axis angle, area and perimeter were then generated.

#### b. Quantitative descriptors of cell shape and migration within toolbox

From the above parameters, the required metrics within the tool box was constructed as indicated in Figure 1. The displacement vector for each frame was calculated by the distance travelled by the centroid from preceding (*n-1*) frame to current (*n*) frame. Global turning angle (GTA) (θ) was calculated as the angle made by displacement vector with the X-axis. Relative turning angle (α) was calculated between current displacement vector (between *n-1* and *n* frames) and successive displacement vector (between *n* and *n+1* frames). Major axis (MA) endpoints were calculated from the length and centroid values generated before. The morpho-migratory angle (μ) was obtained by calculating the acute angle between major axis and the successive displacement vector (between *n* and *n+1* frames). The MA dynamics (Φ) was obtained by calculating the difference in MA angle (generated earlier) in current and preceding frame. Elongation (ε) was calculated by the ratio of minor axis length to major axis length. The persistence ratio was calculated as the ratio between Euclidean distance and accumulated distance.

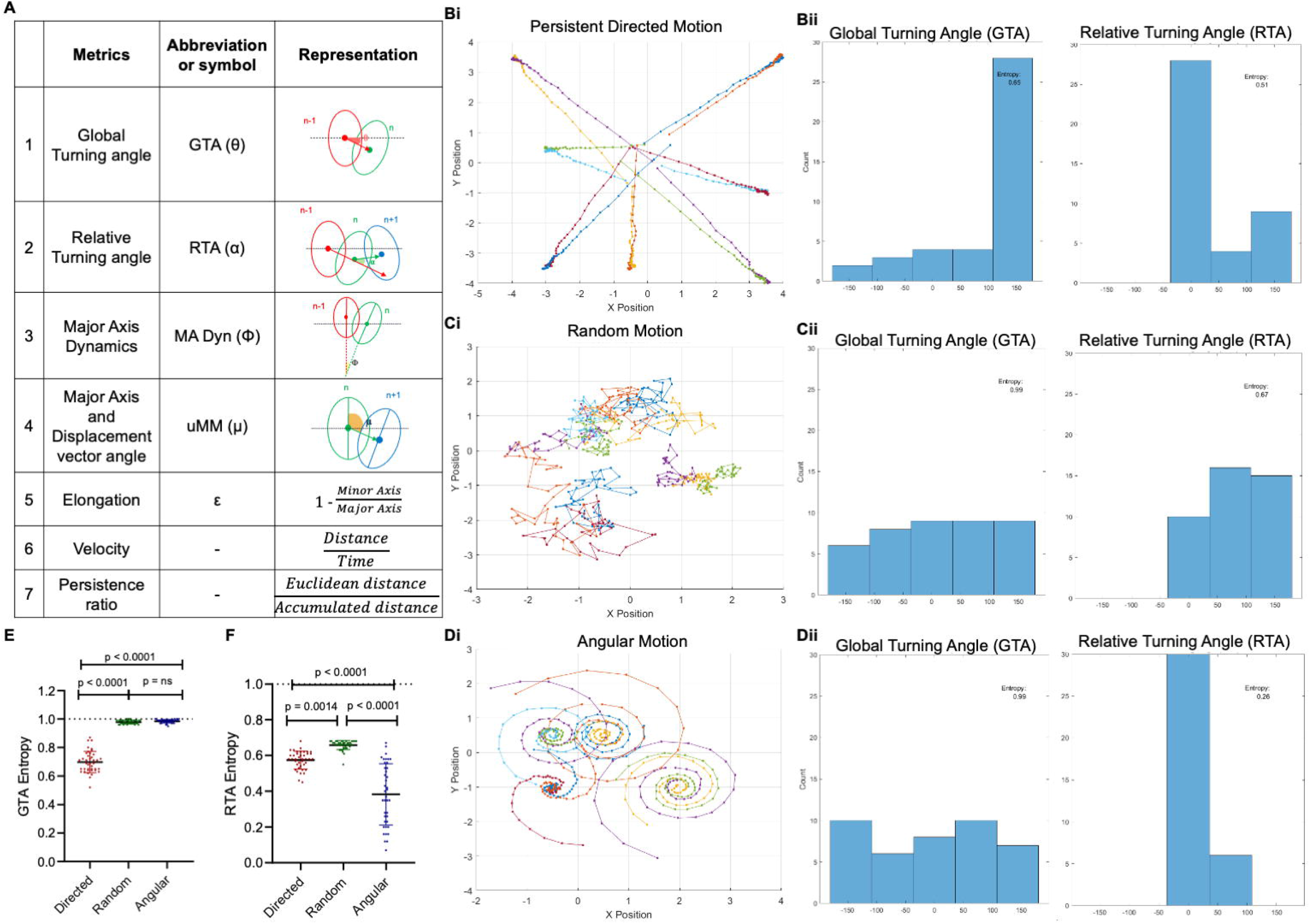

Further, mean squared displacement (MSD) was calculated using the formula

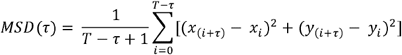

where MSD(τ) represents the MSD for a single cell with step size τ in a total time of T. Since, the frame acquisition rate was every 2 minutes, step size was increased in increments of 2 up to 60. To obtain a population average for each step size, MSDs were averaged over all cells. Log of average MSD and step size was calculated and plotted to fit a linear regression curve and obtain the slope value (k). Similarly, root mean squared (RMS) of turning angle and elongation metrics was calculated with an additional square root calculation of the mean squared values. All the parameters were calculated using MATLAB.

### 4. Statistical Analysis

All time-lapse experiments were performed thrice or more (mentioned otherwise). GraphPad Prism 8.0.1 was used for generating graphs and statistical analysis. For statistical analysis, one-way ANOVA was performed. The correlation matrix (seen in Figures 5E and F) is performed using a corrplot package in R, based on Pearson’s correlation coefficient, between all biological replicates, after which the average of the values is considered. The MATLAB codes used in the above study are available in the following github link: https://github.com/madhumitha-rsuresh/Morphomigratory-parameters

## RESULTS

### 1. Integration of a morpho-migration toolkit with entropic measurements reveals distinct modes of migration

The metrics we have calculated in the context of invasive ovarian cancer cells in this manuscript were chosen to estimate cell motility, shape, orientation, and derive their correlations with their migratory direction. These metrics and their representative formulae or pictorial representations are shown in Figure 1A. To elucidate the ‘uncertainty’ in cellular state change over time, Shannon entropy (H) was used to decode the metric flux. Shannon entropy (H) was calculated using the formula,

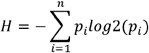

where *n* represents the total number of events and *p*_*i*_ represents the probability for each event (27). Using this definition, entropy for each metric within an individual cell was calculated by sorting every metric value within the given time frame into specific ranged bins and measuring the probability for each bin. Here, n represents number of bins and p represents the probability of a given metric found within the bin at specific time-point. Hence, this gives us a clear histogram depicting the probability distribution of the corresponding metric values within the bins. Furthermore, the entropy values were normalized with the maximum possible entropy (H), which corresponds to equal probability occurrence in all bins, to obtain a dimensionless quantity from 0 to 1 for easier comparison between different scenarios. The Shannon entropy used here goes with the conventional Gibb’s entropy interpretation that an entropic value closer to unity corresponds to ‘highly random’ dynamics while a value closer to ‘0’ corresponds to ‘ordered dynamics’.

To test the sensitivity of the toolkit, synthetic data corresponding to specific migratory modes was generated using a modified Ornstein-Uhlenbeck equation (28) involving drift, diffusion, and an additional rotational term.

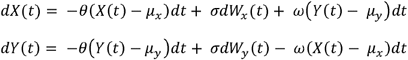

Here, *θ* denotes the rate of reversion to the mean; μ is the mean position; *σ* denotes the volatility of the process (corresponds to random fluctuation intensity); ⍰ denotes rotational frequency; *dW*_*t*_ denotes Wiener process (Brownian motion). X and Y correspond to centroid positions that evolve over time under the above equations. Further, the above equations were discretized for simplification using a constant step size (Δt) of 0.25 for a total time frame of 10 mins.

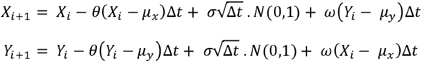

Here, *N(0,1)* denotes a standard normal random variable. Figure 1Bi, Ci, and Di denotes three different migratory modes, namely, persistent directed, random, and angular motion, obtained via tuning the values of θ, μ, σ, and ⍰. For persistent directed motion, the values of μ were kept 3 to 4 coordinates away from the initial point with minimal fluctuation (σ = 0.05) and null rotational frequency. For random motion, the μ values were kept closer to the initial starting point with a higher fluctuation rate (σ = up to 0.5) and null rotational frequency. For angular motion, the values of μ and ⍰ (⍰ = up to 1.5) were allowed to vary, but the fluctuation rate was kept minimal (σ = 0.05).

The synthetic tracks generated using the above equation for different migratory modes are shown in Figure 1B-D. The variation in entropy values for distinct migratory metrics, specifically global and relative turning angles (GTA and RTA, respectively), clearly indicated the type of migration the particles have undertaken (Figure 1E and F; graph depicting mean entropies of GTA and RTA, respectively, for particles of different migratory modes; statistical analysis is done using one-way ANOVA). While the global turning angle (GTA) corresponds to the current direction in which the particle has moved with respect to a fixed axis, the relative turning angle (RTA) corresponds to the trajectory the particle has taken with respect to its previous position (29). Figures 1E and F shows a comparison between the mean entropy for the two metrics for individual trajectories. It was seen that the particles that move in a persistent directed motion have lower mean GTA and RTA entropy (angle distributions shown in Figure 1Bii; see Figure 1E and F). In comparison, those having a random/Brownian motion have higher mean GTA and RTA entropy (angle distributions shown in Figure 1Cii see Figure 1E and F). In the case of angular motion, a constant small turn results in lower mean RTA entropy, while the mean GTA entropy increases due to a wider directional range (angle distributions shown in Figure 1Dii; see Figure 1E and 1F). Hence, the synthetic data confirms the efficacy of the toolkit to differentiate between specific migratory modes based on direction and persistence.

### 2. Substrata stiffness influences the rate of migration and persistence of ovarian cancer cells

To compare the impact of the biophysical cues of the substrata matrix microenvironment on the migratory dynamics of ovarian cancer cells, polyacrylamide (PA) hydrogels of Young’s modulus 0.5 kPa (resembling stiffness seen in non-cancerous glandular tissues) and 20 kPa (seen in fibrosed areas of tumors such as the omental cake in epithelial ovarian cancer metastasis) were used. Both SK-OV-3, a mesenchymal aggressive cell line and OVCAR-3, an epithelioid metastatic cell line, when cultivated on soft substrata, moved with a slower velocity than those on stiffer substrata (Figure 2A-D; images represent migration trajectories with a color heatmap representing velocity values at different time points; inset of a single trajectory shown on the right). Confirmation of lower mean velocity values of low stiffness-cultured cells is also shown in Figure 2E (graph depicting individual cell velocities on soft and stiff substrata; significance computed using one-way ANOVA). Interestingly, OVCAR-3 cells, known to have greater epithelioid characteristics, were faster than mesenchymal SKOV3 cells agnostic of stiffness. Furthermore, the mean entropy for velocity was also higher for both cells on stiffer substrata compared to softer substrata, as shown in Figure 2F (graph depicting mean entropies of cell velocity on soft and stiff substrata; significance computed using one-way ANOVA) as well as in Figure 2C and D, where the heatmap reveals motion driven by constant switching between low and high velocity rates compared with Figure 2A and B.

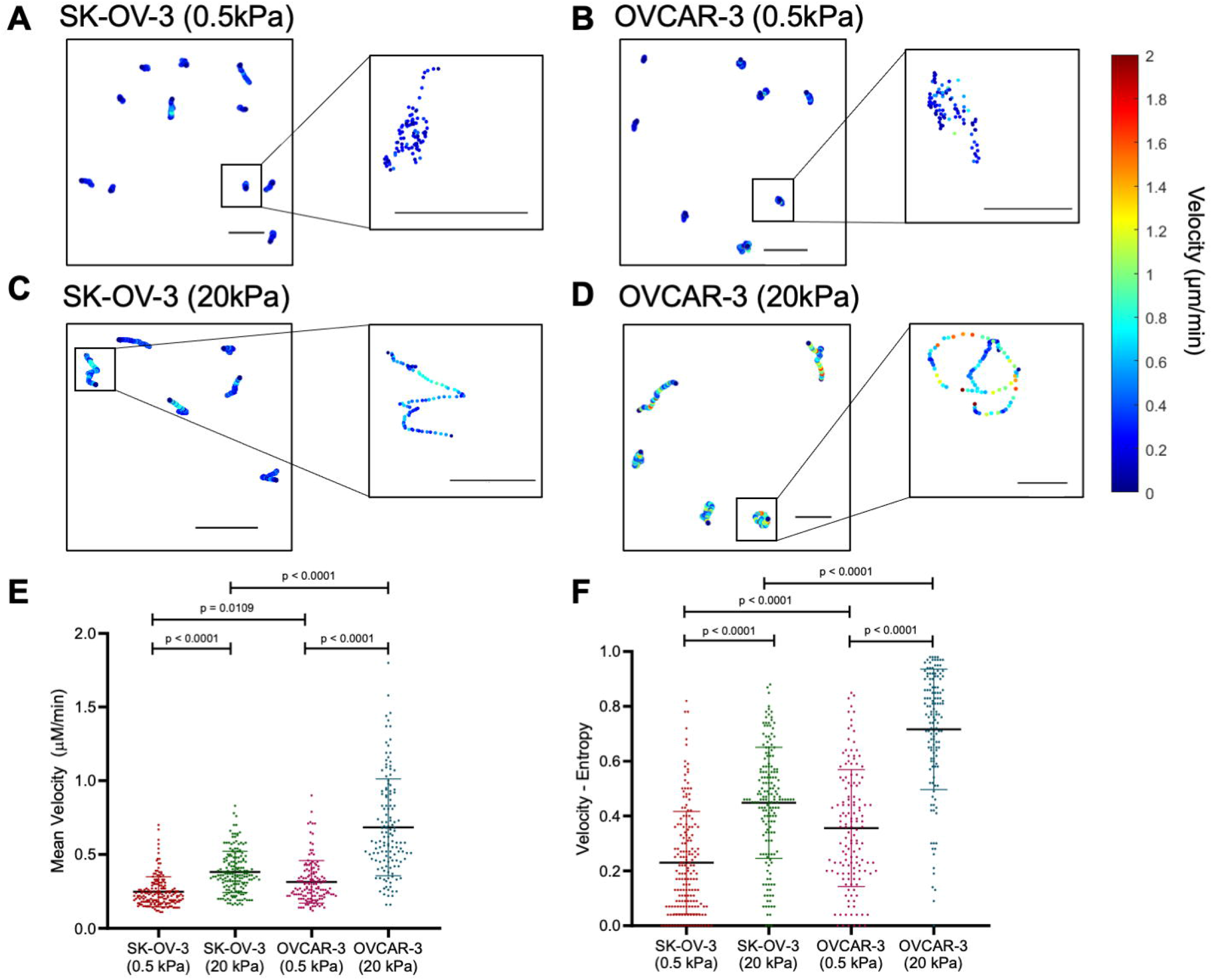

The substrata-contextual difference in migration velocity led us to ask whether OVCAR-3 and SK-OV-3 cells have distinct migration modes, especially in stiffer matrix microenvironments. The possibility of distinct migratory modes was initially characterized by measuring mean squared displacement (MSD) values as a function of time interval τ, fit into a power law to identify the diffusivity of the system. A constant slope value of k=1, indicates random motion while that of k=2, indicates ballistic motion. Other values, such as those between 0 to 1 indicate sub-diffusive motion while those between 1 to 2 indicate superdiffusive motion (30). Both SK-OV-3 and OVCAR-3 cell types showed superdiffusive motility on low and high stiffness as shown in Figure 3E. However, the migration of OVCAR-3 cells on lower stiffness was more random (k = 1.07) when compared with on higher stiffness (k = 1.44). SK-OV-3 cells, on the other hand, showed directed motion in both cases, with increasing stiffness increasing the directionality further (k = 1.58 and 1.78 for 0.5 kPa and 20 kPa gels, respectively). This was further quantified and interpreted through measuring the global turning angles (GTA) and relative turning angles (RTA) (Figure 3A-D, where A and B represent the time-resolved migration in the centroids of representative SK-OV-3 cells on 0.5 kPa and 20 kPa gels respectively and C and D represent the same for OVCAR-3 cells).

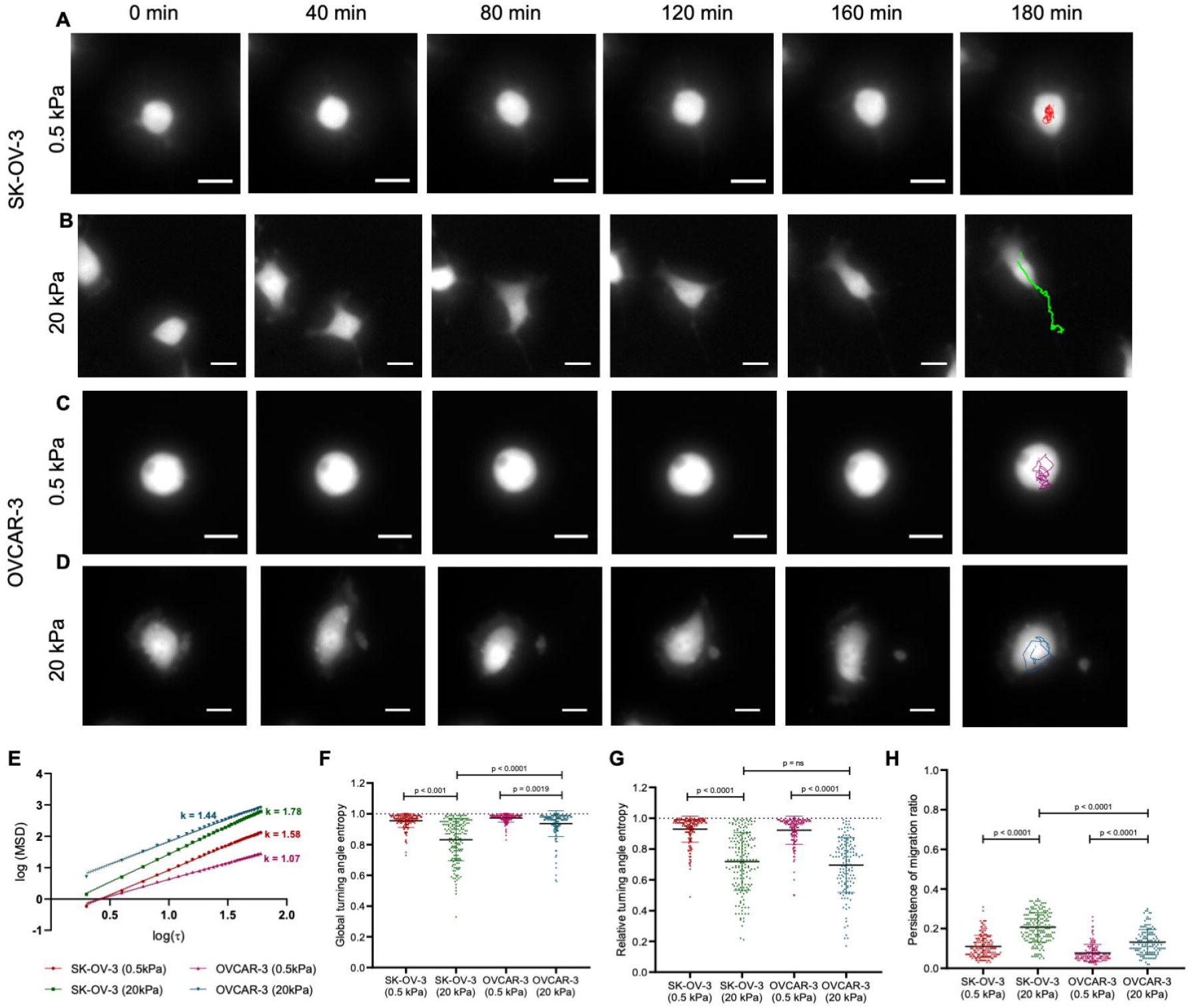

The mean GTA entropy for both cells on lower stiffnesses was high, suggesting lower persistence in motion; on stiffer gels, SK-OV-3 had a lower mean GTA than OVCAR-3 indicating a more persistent migration for the former (Figure 3F; significance computed using one-way ANOVA). The mean RTA entropy for both cells on lower stiffnesses was also high; on stiffer substrata, whereas the mean RTA entropy of SK-OV-3 was low, this was (in contrast to GTA entropy) also found to be low for OVCAR-3 cells (Figure 3G; significance computed using one-way ANOVA). Compared with the entropic signatures seen for our synthetic data in Figure 1 and confirmed through the time lapses, SK-OV-3 on high stiffness migrates in a directed manner, whereas OVCAR-3 on high stiffness shows migrates in an angular fashion. This was confirmed through measurement of persistence ratio in SK-OV-3 cells, which showed higher values than OVCAR-3 cells, with both cells showing greater persistence on stiffer substrata than on softer controls (Figure 3H; graph depicting mean persistence ratios on soft and stiff substrata; significance computed using one way ANOVA). The RMS values of GTA for both cells on both stiffness showed time invariant behavior (Figure S1A). While the RMS values of RTA was invariant of SK-OV-3, for OVCAR-3 especially on high stiffness substrata there was a marked increase, which can be explained by the increase in relative turn calculated when the time scales for measurement in angular motion are lengthened, consistent with the inferred migratory dynamics seen in Figure 3.

### 3. Ovarian cancer cells show morphology-specific correlation between axes of shape and migration on high stiffness

Thus far, our measurements for assessment of migration explored the spatial dynamics of the centroid of cells. To explore associations between the axes of their polar shapes and their motion, the change in major axis orientation (Φ) between consecutive time frames was measured over time (see Figure 4A and B for time-resolved representative photomicrographs of SK-OV-3 and OVCAR-3 cells on stiff substrates with major axes determined across 40 min time intervals). Both cell types on lower stiffness showed a wide range of major axis orientation entropies (Figure 4C; graph depicting mean entropies of major axis orientation on soft and stiff substrata; statistical analysis was performed using one-way ANOVA). However, this might arise due to the more rounded morphology of cells in lower stiffness, resulting in random ellipse and major axis fits disallowing us from inferencing any biological significance to these observations. Interestingly, on higher stiffness, the change in orientation drastically differed between OVCAR-3 and SK-OV-3: the latter, which is found to have persistent directed motion shows relatively lower variation in axis orientation dynamics than OVCAR-3 indicating their ability to polarize and move in the same direction. OVCAR-3 cells, on the other hand, have very high Φ values on stiff substrata, indicating a constant change in orientation during its migration, which could be explained by its angular trajectories indicated in Figure 3.

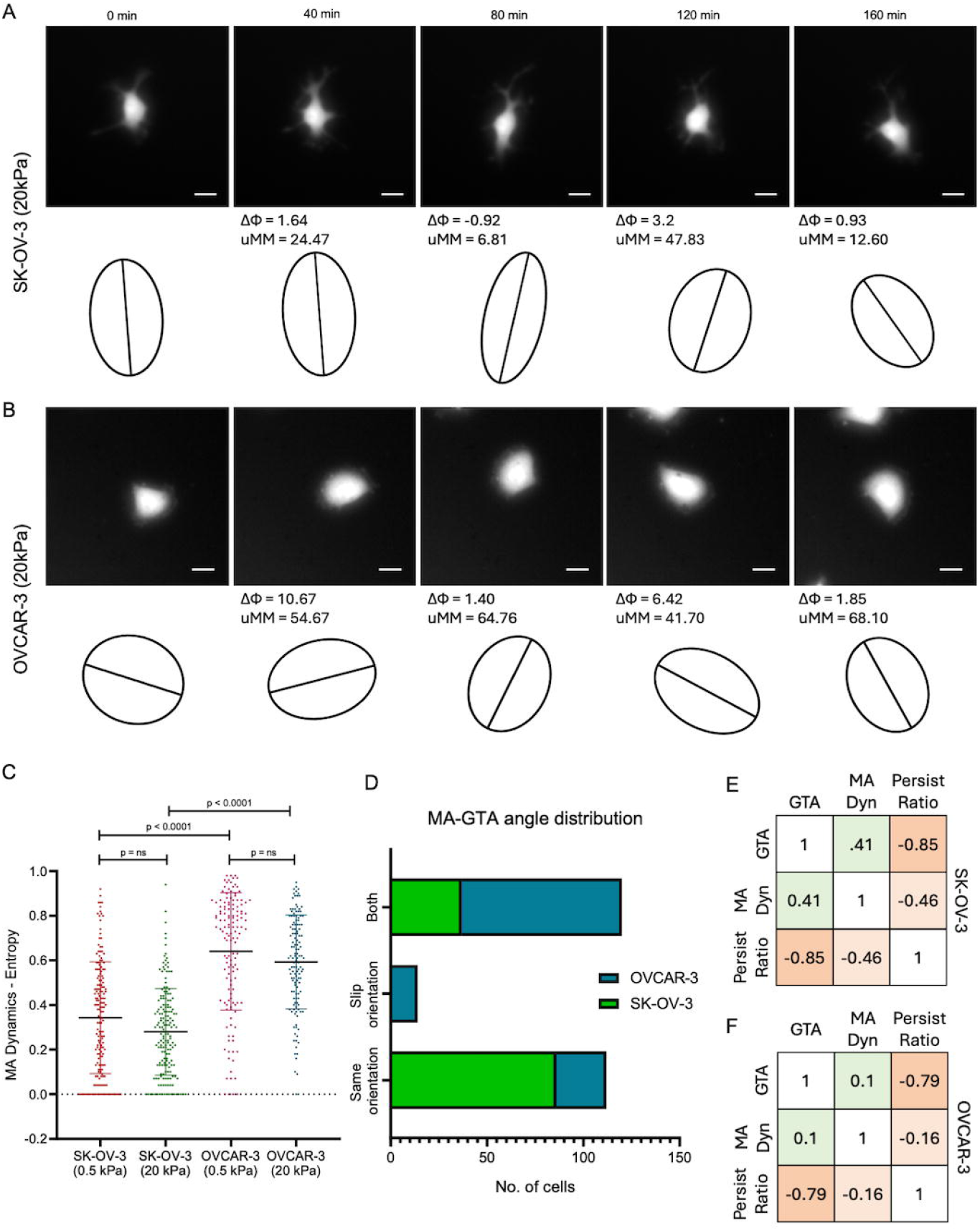

To extrapolate this finding and correlate it with orientation and migration, the uMM angle was measured for each cell (see Figure 1A for definition). A wide range of SK-OV-3 cells had their trajectory with a very low uMM angle (<45°). This confirms that the minimal orientation dynamics are due to their ability to polarize and move in that direction. One intriguing phenomenon we found in OVCAR-3 cells was that most cells had uMM angles greater than 45° as seen in Figure 4A and B. This denotes that the direction in which cells move is unrelated to the current axial orientation they currently exhibit. If a cell shows such a high uMM angle for more than 60% of its time frame, we refer to such cells as exhibiting ‘slip motion’; Figure 4D shows that the phenomenon was predominantly observed only in OVCAR-3. A correlation matrix between entropic values of GTA, and MA dynamics, and persistence ratio for both the cell types also revealed relatively greater correlation between migratory direction and the orientation dynamics in SK-OV-3 cells (Figure 4E; correlation = 0.41 between entropic values for MA dynamics and GTA entropy) than OVCAR-3 cells (Figure 4F; correlation = 0.1). This gives a new insight into understanding the impact of high stiffness on the migratory capacity of epithelioid cancer cells as they undergo a constant velocity and polarity change (often uncorrelated) to push through different directions.

### 4. Global deformability of cells increases with an increase in stiffness

We next asked whether ovarian cancer cells on higher stiffness underwent morphological axis alteration through a global shape alteration. To address this, we measured the elongation ratio (see Figure 5A) from best-fitted ellipse parameters and calculated its distribution to measure elongation entropy values. As seen in Figure 5A-D (which shows time-resolved shape changes in SK-OV-3 and OVCAR-3 cells grown on softer and stiffer substrata respectively, with their global cell shapes denoted using distinct colors), both cell types on lower stiffness were less deformable, when compared to on higher stiffness, suggesting that the cells behaved like rigid objects with a rounded morphology, as indicated by lower mean elongation values (Figure 5E; graph depicting mean values of elongation ratio of cells on soft and stiff substrata; statistical analysis was performed using one-way ANOVA). Both cells also showed high entropic values for elongation dynamics on high stiffness compared with on softer substrata, as seen in Figure 5F (graph depicting mean entropy of elongation ratio of cells on soft and stiff substrata; statistical analysis is done using one-way ANOVA), indicating the significance of deformability for migration. Further, the RMS values of elongation showed time step invariance (Figure 5G), indicating the dynamicity to be held true for over shorter and longer time intervals. Although, on stiffer environments, the entropy between the two cell types was insignificantly altered, the fold change in deformability for OVCAR-3 across stiffnesses was higher suggesting a more substrate pliable cytoskeleton that could explain its higher exploratory migratory behavior through angular and slip locomotion.

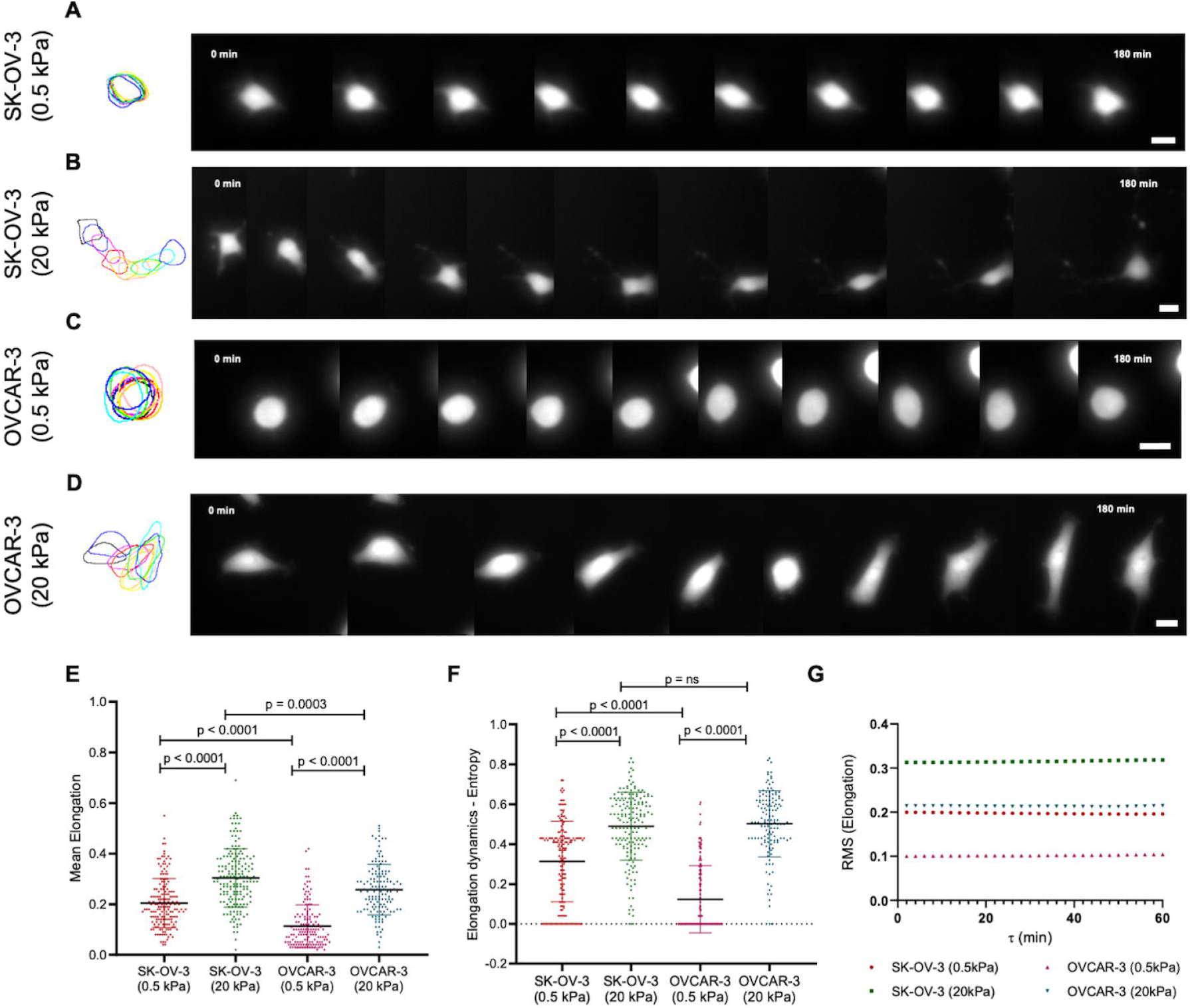

## DISCUSSION

In this study, we propose a simple yet highly sensitive approach to capture the spatiotemporal dynamics of cell morphology and migration by combining mathematical imaging metrics and the concept of Shannon entropy. The metrics used here involve common descriptors, such as velocity, turning angles, persistence ratio, and elongation, as well as recently developed metrics, such as major axis dynamics and morpho-migrational angle (17). This gives us insights into cell trajectory, direction, orientation, and geometry over time. Recent studies by Liu and coworkers, have shown the significance of calculating Shannon entropy for being able to capture dynamicity of, and distinguish between, distinct cellular migratory modes, such as persistence or random walk, and morphological features, such as amoeboid and mesenchymal motility (31–33). Hence, a pipeline seeking to integrate both sets into a single framework with an effort to seek correlative links between the two can help bridging the gap between intuitive verbal interpretations and construction of complex theoretical models to quantify cellular plasticity.

In this study, ovarian cancer cells were found to be highly sensitive to biochemical and biophysical cues from their microenvironment, resulting in distinct phenotypic behaviors. Tuning the stiffness of the underlying substratum, while keeping the ligand concentration and structure same, gave deeper insight into the effect of elastic regime of the substrata environment on cell deformability and motility. Current studies on understanding the role of substrate stiffness on ovarian cancer progression have looked majorly at static quantities but they held true with the conventional interpretation that cells on higher stiffness have higher migratory capacity and spreading (34). However, our analysis shows that compared to the consistent but unsurprising directed persistent behavior of the mesenchymal SK-OV-3, the migratory mode of epithelioid OVCAR-3 cells on higher stiffness showed novel behaviors (Figure 4). Epithelioid OVCAR-3 cells were in fact seen to surpass SK-OV-3 cells in their mean velocity distribution on higher stiffness as well as angular and slip motility indicating a more exploratory behavior that would be possible on fibrosed tissues within the peritoneum. Our observations underscore the need to move beyond a conventional narrative of a relative sessility and motility as shown by an epithelial and a mesenchymal cell type.

Recent studies have been investigating the interconvertibility of epithelial, mesenchymal, and amoeboid cell states (8). Mesenchymal motility is known to be possible through filopodial or lamellipodial formation, where cell-matrix interaction plays a major role in the translocation of the cell. The mesenchymal cells are known to exert force on the matrix, which causes a turnover pulling from the matrix that pushes the cell forward through actomyosin contractility. Amoeboid cells, on the other hand, achieve much higher velocity rates than mesenchymal cells by forming blebs, actin-rich pseudopodia, or highly contractile uropods. Their ability to ‘shape-shift’ is attributed to constant reorganization of their actin cytoskeleton through rapid recycling (35). The ability of OVCAR-3 cells to show higher mean velocity than mesenchymal SK-OV-3 suggests that OVCAR-3 cells might indeed display amoeboid characteristics on higher stiffness. Recent investigations on the ability of cells to switch between different cellular states has indicated that amoeboid state might lie within a larger spectra of state transitions within which epithelial and mesenchymal states reside (7,36). Furthermore, the ability of cells to reside in a hybrid state is consistent with the complexity and heterogeneity observed in cancer cell populations (37). In our study, the reduction in persistence ratio (Figure 3H) and increase in angular motion (Figures 3F and G) seen in OVCAR-3 cells at high stiffness, suggests that they might exhibit hybrid migrational behavior, as they lose their directionality but continue to show higher cellular velocity (Figure 2D). Representative snapshots of time-lapse images of OVCAR-3 cells on high stiffness (Figure 3D and 4B), also show protrusion of fan-like lamellipodia during migration, which resembles actin polymerization-driven amoeboid structures (38). SK-OV-3 cells, on the other hand, are seen to show an elevated mesenchymal character with increased persistent directed motion on high stiffness.

Aberrant cell polarity signaling has also been known to be the leading cause of EMT, which in turn causes higher cancer invasiveness and metastasis (39). The loss of cellular polarity with respect to the direction of migration was also highly seen in OVCAR-3 cells on high stiffness, where cells moved in a direction unrelated to the orientation of the cells. This loss in correlation between orientation and direction of migration, which has been referred to as slip motion, was seen to be a unique trait taken up by OVCAR-3 cells on high stiffness (Figure 4B and F) and resulted in a constant change in their orientation as they underwent consistent tilting during angular motion. This was quantified and confirmed by high entropic values of MA dynamics for OVCAR-3 cells on high stiffness (Figure 4C). Such hybrid behavior suggests that OVCAR-3, in general, might have higher mechanosensitivity and adaptive capacity to switch between different transition states, to increase its migratory and invasive potential. On the other hand, SK-OV-3 cells were found to have typical mesenchymal characteristics by forming lamellipodia or filopodia in the direction of motion, retaining cellular orientation in the direction of motion (Figure 4A and E). Stiffness was also seen to induce higher global deformability in both the cell types, indicating that morphodynamics and the microenvironment may cooperate to facilitate cells to attain their specific migratory mode (Figure 5).

In summary, we demonstrate that a rigorous imaging approach combining morphology and migration metrics helps probe the spatiotemporal dynamics of the cellular phenotype, which can be extensively used to study pathological states such as cancer. In addition, it is seen that an interplay between the mechanical microenvironment and morphological traits produces distinct migratory modes in different ovarian cancer sub-types. While SK-OV-3 cells show elevated mesenchymal characteristics on high-stiffness gels, in the same environments OVCAR-3 cells show stunning diversity in migratory exploration and higher velocities. In future studies, we will extend the study to a greater diversity of cell lines and microenvironmental contexts. In addition, we will investigate how these matrix-driven cues will modulate the dynamical phenotype of multicellular cancer collectives.

## Acknowledgments

This work was supported by India Alliance DBT Wellcome Trust Fellowship (IA/I/17/2/503312) awarded to R.B. It was also supported by the John Templeton Foundation (no. 62220), the Indo-French Centre for the Promotion of Advanced Research (CEFIPRA grant: 69T08-2) and the International Emerging Actions (328003) to RB. MS acknowledges the Wells Fargo MTech Women’s Fellowship for support. We thank Claire Valotteau and Felix Rico (INSERM and Aix-Marseille Université) for their inputs on time scale-based dynamical analysis. The opinions expressed in this paper are those of the authors and not those of the John Templeton Foundation.

