## Supplementary File for "Ovarian cancer cells exhibit diverse migration strategies on stiff collagenous substrata"

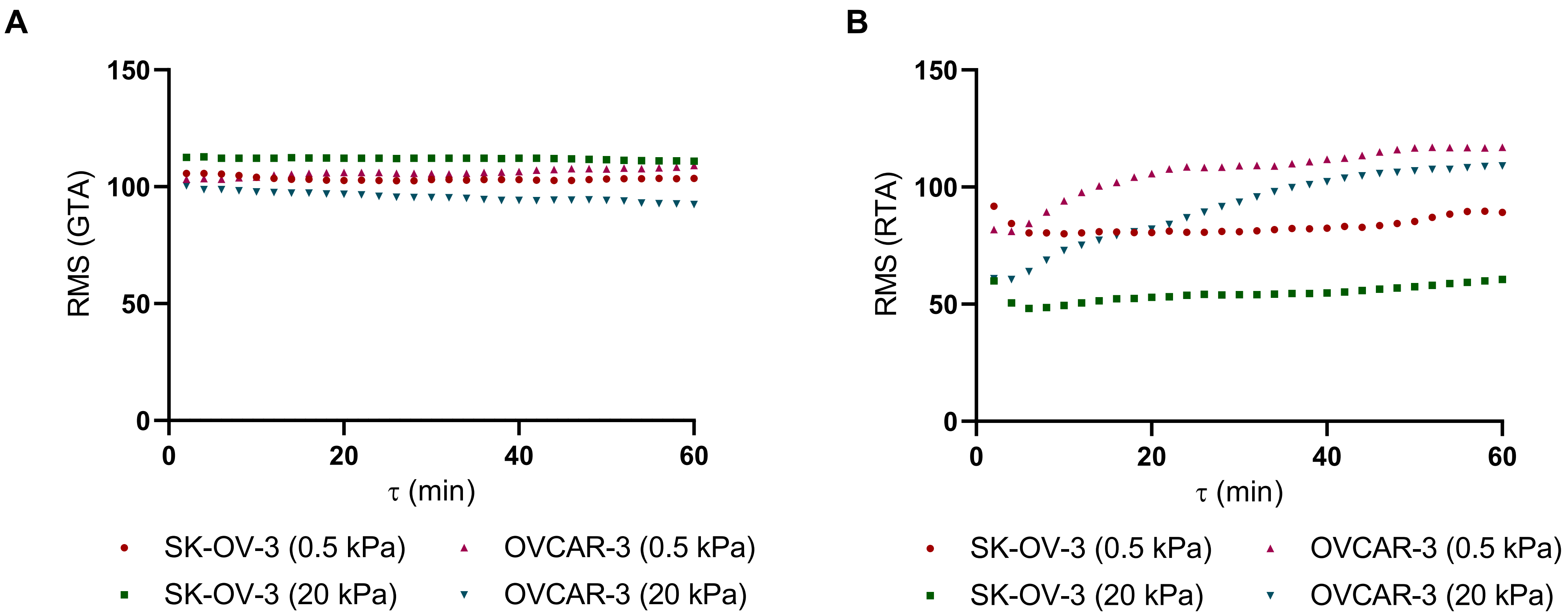


Figure S1: Root mean square values of Global Turning Angle (GTA) (A) and Relative Turning Angle (B) measured across different time scales for SK-OV-3 and OVCAR-3 cells cultured on 0.5 kPa (soft) and 20 kPa (stiff) Collagen I substrata

**Supplementary Figure 1:** Time-lapse video of SK-OV-3 cells on low stiffness (0.5 kPa) for a duration of 3 hours with snapshots taken at a regular interval of 2 minutes (Scale bar: 20 μm)

**Supplementary Figure 2:** Time-lapse video of SK-OV-3 cells on high stiffness (20 kPa) for a duration of 3 hours with snapshots taken at a regular interval of 2 minutes (Scale bar: 20 μm)

**Supplementary Figure 3:** Time-lapse video of OVCAR-3 cells on low stiffness (0.5 kPa) for a duration of 3 hours with snapshots taken at a regular interval of 2 minutes (Scale bar: 20 μm)

**Supplementary Figure 4:** Time-lapse video of OVCAR-3 cells on high stiffness (20 kPa) for a duration of 3 hours with snapshots taken at a regular interval of 2 minutes (Scale bar: 20 μm)

**Supplementary Figure 5:** Time-lapse video of SK-OV-3 cells on high stiffness (20 kPa) for a duration of 3 hours with snapshots taken at a regular interval of 2 minutes (Scale bar: 20 μm)

**Supplementary Figure 6:** Time-lapse video of OVCAR-3 cells on high stiffness (20 kPa) for a duration of 3 hours with snapshots taken at a regular interval of 2 minutes (Scale bar: 20 μm)

**Supplementary Figure 7:** Time-lapse video of SK-OV-3 cells on low stiffness (0.5 kPa) for a duration of 3 hours with snapshots taken at a regular interval of 2 minutes (Scale bar: 20 μm)

**Supplementary Figure 8:** Time-lapse video of SK-OV-3 cells on high stiffness (20 kPa) for a duration of 3 hours with snapshots taken at a regular interval of 2 minutes (Scale bar: 20 μm)

**Supplementary Figure 9:** Time-lapse video of OVCAR-3 cells on low stiffness (0.5 kPa) for a duration of 3 hours with snapshots taken at a regular interval of 2 minutes (Scale bar: 20 μm)

**Supplementary Figure 10:** Time-lapse video of OVCAR-3 cells on high stiffness (20 kPa) for a duration of 3 hours with snapshots taken at a regular interval of 2 minutes (Scale bar: 20 μm)
